# The accelerated infectious disease risk in the Anthropocene: more outbreaks and wider global spread

**DOI:** 10.1101/2020.04.20.049866

**Authors:** Serge Morand, Bruno A. Walther

## Abstract

The greatly accelerated economic growth during the Anthropocene has resulted in astonishing improvements in many aspects of human well-being, but has also caused the acceleration of risks, such as the interlinked biodiversity and climate crisis. Here, we report on another risk: the accelerated infectious disease risk associated with the number and geographic spread of human infectious disease outbreaks. Using the most complete, reliable, and up-to-date database on human infectious disease outbreaks (GIDEON), we show that the number of disease outbreaks, the number of diseases involved in these outbreaks, and the number of countries affected have increased during the entire Anthropocene. Furthermore, the spatial distribution of these outbreaks is becoming more globalized in the sense that the overall modularity of the disease networks across the globe has decreased, meaning disease outbreaks have become increasingly pandemic in their nature. This decrease in modularity is associated with tremendous increases in mobility, especially in air traffic. We also show that those countries and regions which are most central within these disease networks tend to be countries and global regions with higher GDPs. Therefore, one cost of greater economic growth and the associated increased global mobility is the increased risk of disease outbreaks and their wider spread. Finally, the recent global outbreaks of Covid-19 and monkeypox allowed us to demonstrate that the time of first occurrence in each country was correlated with each country’s centrality value in the disease network. We briefly discuss three different scenarios of how mobility may develop in the future which decision-makers might discuss in light of our results.

## Introduction

The Anthropocene has also been nicknamed the ‘Great Acceleration’ because various socioeconomic and Earth-System related indicators experienced a continuous and often exponential growth after the Second World War^1-3^. While this relentless growth of the human enterprise improved human well-being around the world in many aspects^4-6^, negative impacts have likewise increased^7,8^. In tandem, warnings about a climate emergency^9,10^, a sixth mass extinction^11-13^, increasing ocean acidification and dead zones^14,15^, and even widespread ecosystem collapse^12,16,17^ and transgression of safe planetary boundaries^18^ have grown increasingly urgent. To avoid or at least dampen the anticipated or already realized changes, scientists and many others have called out for radical changes to how human economies operate and relate to human societies and their environments (e.g., references 10, 19-23).

We here want to draw attention to another important and noteworthy feature of the Anthropocene which greatly affects public health, human well-being, and economic performance. These findings are especially pertinent as the world reels from the health, social, and economic impact of the current SARS-CoV-2 pandemic (from hereon called Covid-19 for simplicity; references 24-26).

The increasing connectivity of human populations due to international trade and travel^27,28^, the rapid growth of the global transport of wild and domesticated animals^29-32^, and other factors such as the increasing encroachment of human populations on hitherto isolated wild animal populations through loss and fragmentation of wild habitats^33-36^ have led to a great acceleration of infectious disease risks, e.g., the increase in emerging infectious diseases and drug-resistant microbes since 1940^37^, the increase in the number of disease outbreaks since 1980^38^, and the increase in the number of animal disease outbreaks since 2005^39^.

Many studies have suggested possible reasons for this increase. Jones et al.^37^ related the emergence of infectious diseases to human population density and growth, latitude, and wildlife host richness, and Plowright et al.^40^ specified ecological drivers for several emerging infectious diseases. There may be several reasons why the number of outbreaks is increasing. Some have to do with local conditions and factors at the source of the outbreak^41,42^, while others are more regional or global factors such as the increasing number of drug-resistant microbes, higher contact rates with wildlife, increased livestock and global food production, loss and fragmentation of natural habitats caused by urbanization and agricultural intensification, extreme climate events, and others^31,37,40,43-50^.

All these factors combined drive an increase in the number of disease outbreaks. Smith et al.^38^ demonstrated this for human infectious diseases using a dataset of 12101 outbreaks of 215 human infectious diseases, which showed a significant increase in outbreaks since 1980.

To expand Smith et al.’s^38^ analysis to the beginning of the Anthropocene, we investigated whether the number of disease outbreaks has increased since the Second World War. However, we also asked the important question whether these disease outbreaks have not just increased in number, but also increased their spatial spread across the globe. We hypothesized that, due the increasing connectivity and mobility of humans, the global spatial pattern of infectious disease outbreaks also changed in response. In other words: have the disease outbreaks become more globalized in the sense that these outbreaks are increasingly shared by countries worldwide?

To investigate this hypothesis, we used the most complete, reliable, and up-to-date global dataset 51. This dataset can be used to enumerate the annual number of disease outbreaks. To investigate the changing global patterns of disease outbreaks, we used this dataset to calculate two measures which were recently introduced into ecological and parasitological studies. These two measures, namely modularity and centrality, quantify the connectivity of bipartite networks and have already been used to investigate various ecological, parasitological, and epidemiological questions during the last two decades (e.g., references 52-59).

Modularity is defined as the extent to which nodes (specifically, sites and species for presence-absence matrices) in a compartment are more likely to be connected to each other than to other nodes of the network which are outside of that compartment^60^. The calculation of a modularity measure is useful for global phenomena because it allows the overall level of compartmentalization (or fragmentation) into compartments (also called clusters, modules, subgroups, subsets) of an entire dataset to be quantified. High modularity in a global network means that subgroups of countries and disease outbreaks interact more strongly among themselves (that is, within a compartment) than with the other subgroups (that is, among compartments)^58^. Changes in modularity can also be visualized graphically, and we used three different visualizations, namely bipartite networks, bipartite modules, and unipartite networks.

Centrality is defined as the degree of the connectedness of a node (e.g., a keystone species in ecological studies^53,54^). In the context of our study, centrality is the degree of the connectedness of a country and those countries connected to it. We thus estimated the countries which are the potential centres of disease outbreaks by investigating the eigenvector centrality of a given country in a network of countries which share disease outbreaks among each other. Eigenvector centrality is a generalization of degree centrality, which is the number of connections a country has to other countries in terms of sharing disease outbreaks. Eigenvector centrality considers countries to be highly central if the countries connected to them through shared outbreaks are also connected to many other well-connected countries^61,62^.

Using the widely used global dataset on infectious disease outbreaks^51^, we here present results which demonstrate that the accelerated number of disease outbreaks and their increased global spread are two further threatening aspects of the accelerated infectious disease risk associated with the globalization process which characterizes the Anthropocene.

## Results

### Increase in global measures of mobility and GDP

The increases in measures of mobility during the Anthropocene have been staggering, with growth percentages of up to 5600% (Table 1). Many more impressive local, regional, and historical examples exist (e.g., references 28, 63, 64). For further analyses below, we only used the World Bank data on flight passengers and air freight; therefore, we detail them here. Since the 1970s, the total global number of flight passengers (Fig. 1A) and air freight (Fig. 1B) have increased almost continuously except for 2020 (Supplementary Data S1). The total number of flight passengers increased from 310 million in 1970 to 4 billion and 556 million in 2019, which is an almost 15 times increase. Due to the Covid-19 pandemic, the number of passengers decreased to 1 billion and 809 million in 2020 (about two-fifths of the 2019 level). The total amount of air freight increased from 15569 million MT-km in 1973 to 221495 million MT-km in 2019, which is about a 14 times increase. In 2020, the total amount of air freight also decreased to 18053 million MT-km (less than one-tenth of the 2019 level). Global GDP per capita also increased exponentially from 459 USA dollars per capita in 1970 to 11407 USA dollars per capita in 2019, which is an almost 25 times increase (Fig. 1C). The global GDP per capita also decreased due to the Covid-19 pandemic to 10936 USA dollars per capita in 2020 but this represents only a 4% decrease compared to the 2019 level. Even when we control for the number of the global human population, almost the same growth trends are evident (Fig. 2A-C), meaning that increased global mobility and GDP are driven by global population growth as well as by individual increases in mobility and wealth.

**Fig. 1.**
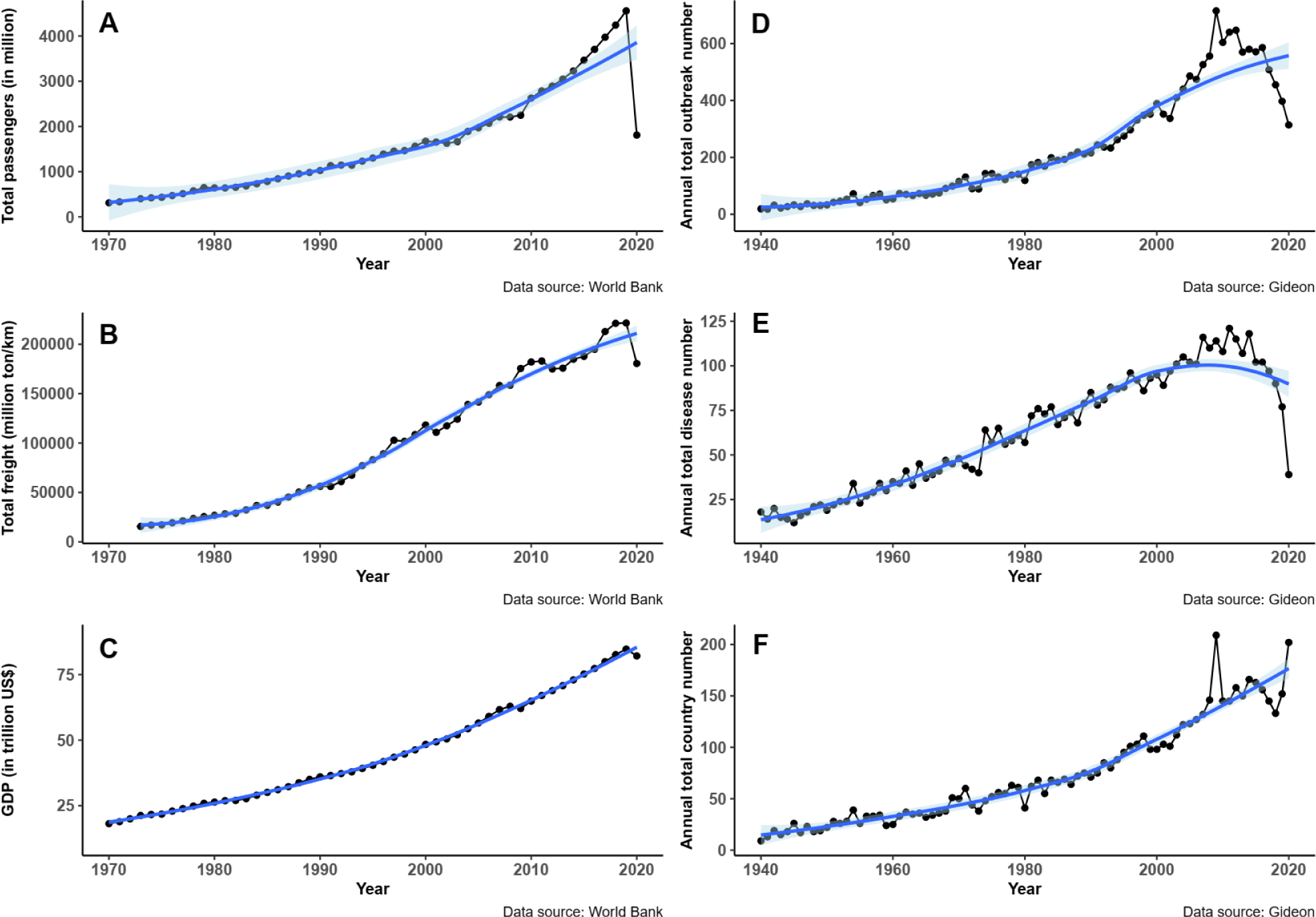
The increase of air travel, GDP, and disease outbreaks since the 1940s. Resulting graphs from plotting the total global number of air passengers from 1970 to 2020 (A), total global air freight from 1972 to 2020 (B), and global GDP from 1970 to 2020 (C); and the annual total outbreak number (D), the annual total disease number (E) and the annual total country number (F) from 1940 to 2020. Fitted smooth regressions (in blue) with confidence intervals (in light blue) are shown.

**Fig. 2.**
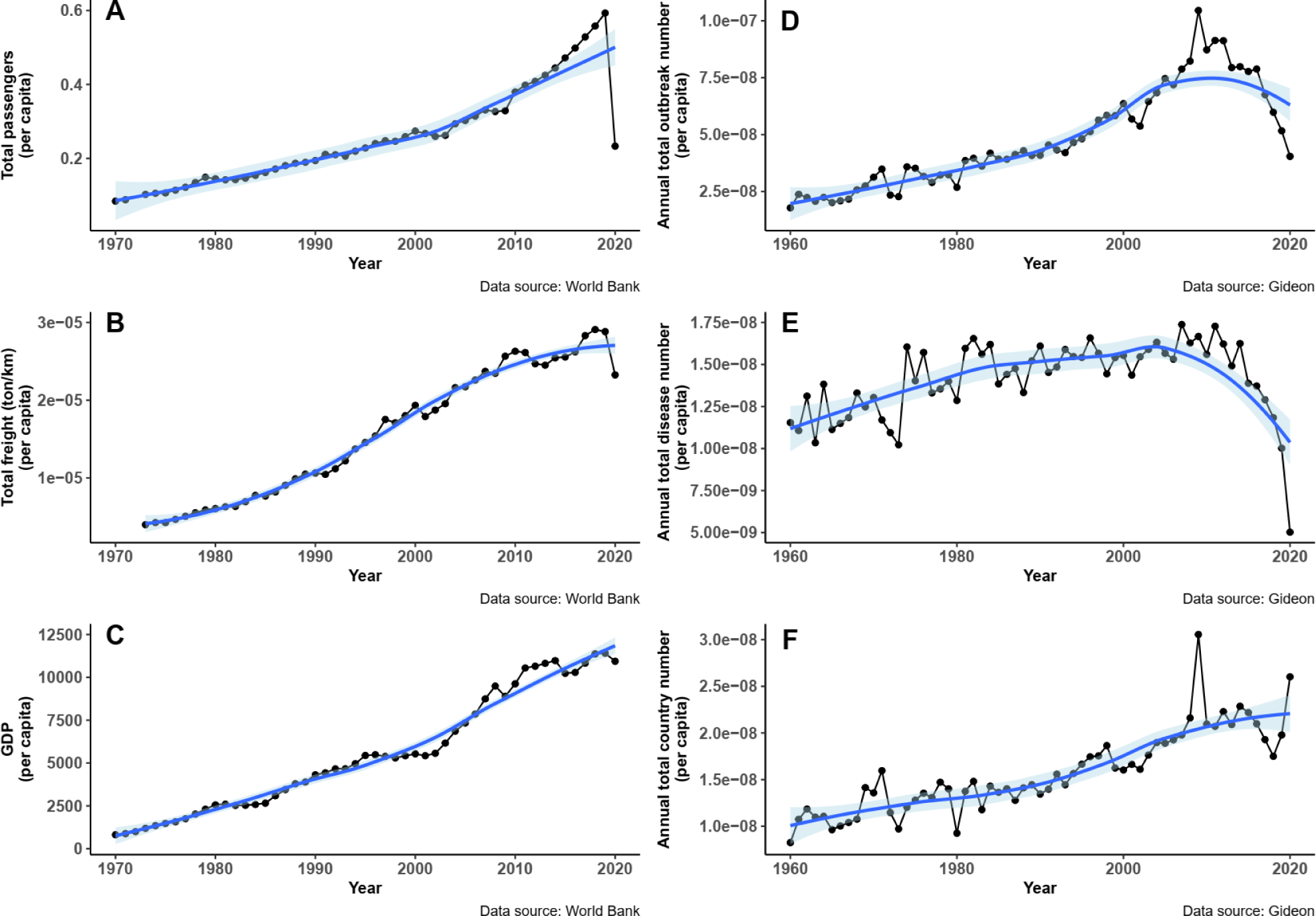
The increase of air travel, GDP, and disease outbreaks since the 1940s controlled by population. Resulting graphs from plotting the same data as in Fig. 1, but controlled for by global human population (per capita). Fitted smooth regressions (in blue) with confidence intervals (in light blue) are shown.

**Table 1.**
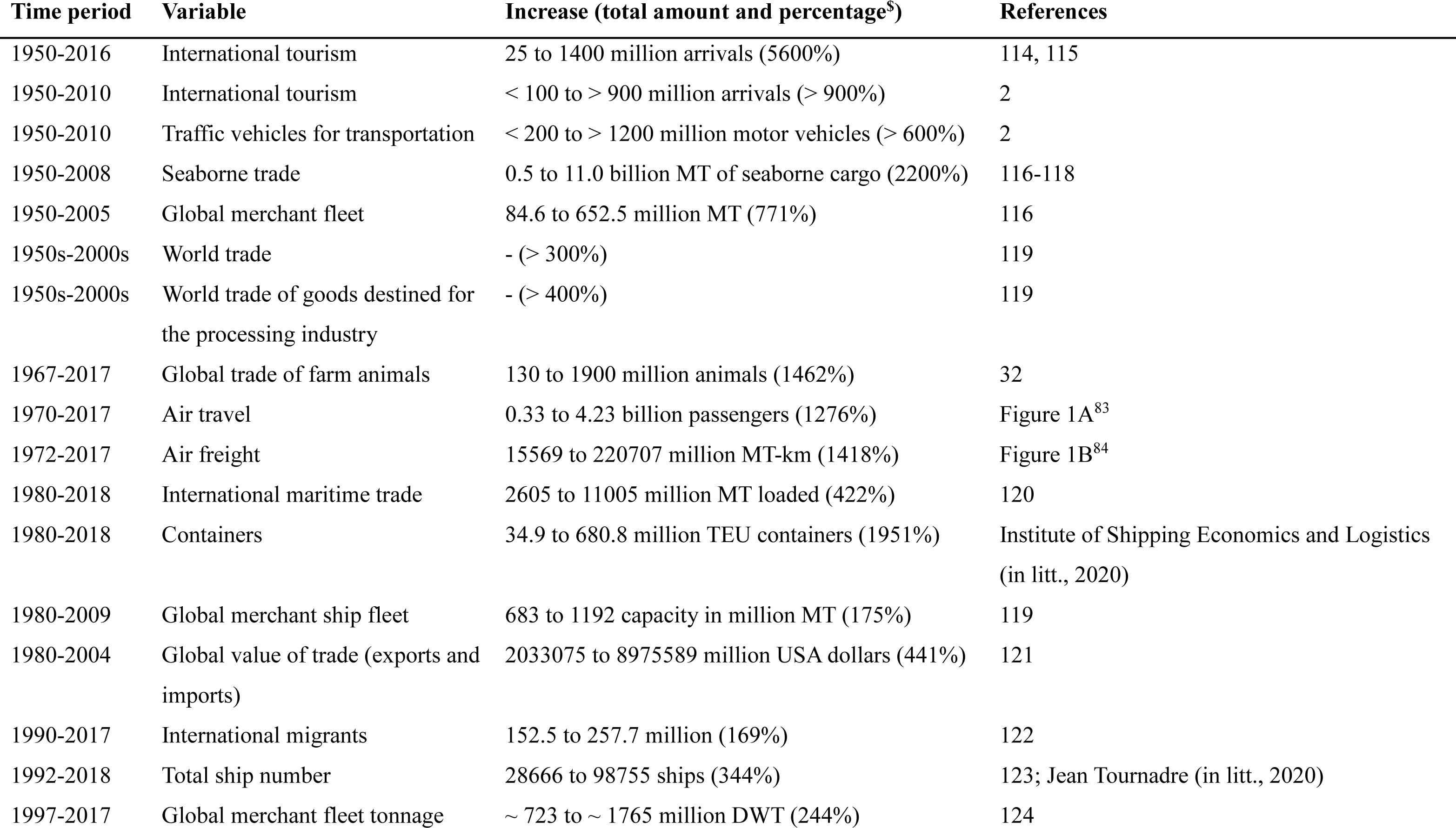
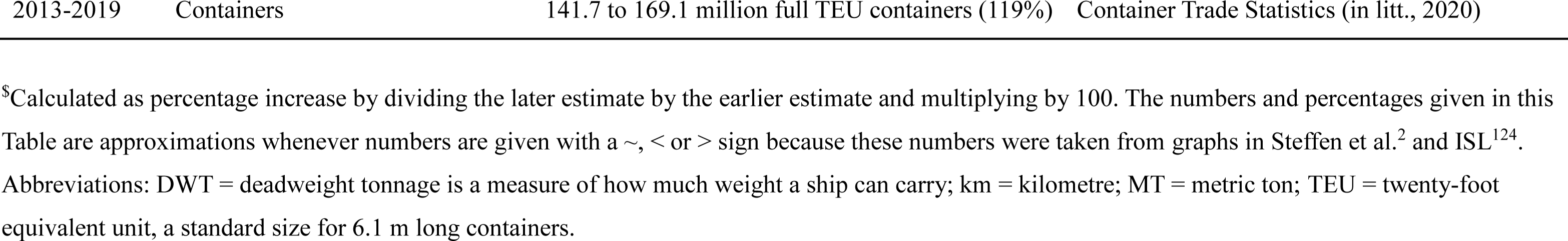
Increase in global measures of mobility during the Anthropocene.

| **Time period** | **Variable** | **Increase (total amount and percentage<sup>s</sup>)** | **References** |
| --- | --- | --- | --- |
| 1950-2016 | International tourism | 25 to 1400 million arrivals (5600%) | 114, 115 |
| 1950-2010 | International tourism | < 100 to > 900 million arrivals (> 900%) | 2 |
| 1950-2010 | Traffic vehicles for transportation | < 200 to > 1200 million motor vehicles (> 600%) | 2 |
| 1950-2008 | Seaborne trade | 0.5 to 11.0 billion MT of seaborne cargo (2200%) | 116-118 |
| 1950-2005 | Global merchant fleet | 84.6 to 652.5 million MT (771%) | 116 |
| 1950s-2000s | World trade | - (> 300%) | 119 |
| 1950s-2000s | World trade of goods destined for the processing industry | - (> 400%) | 119 |
| 1967-2017 | Global trade of farm animals | 130 to 1900 million animals (1462%) | 32 |
| 1970-2017 | Air travel | 0.33 to 4.23 billion passengers (1276%) | Figure 1A<sup>83</sup> |
| 1972-2017 | Air freight | 15569 to 220707 million MT-km (1418%) | Figure 1B<sup>84</sup> |
| 1980-2018 | International maritime trade | 2605 to 11005 million MT loaded (422%) | 120 |
| 1980-2018 | Containers | 34.9 to 680.8 million TEU containers (1951%) | Institute of Shipping Economics and Logistics (in litt., 2020) |
| 1980-2009 | Global merchant ship fleet | 683 to 1192 capacity in million MT (175%) | 119 |
| 1980-2004 | Global value of trade (exports and imports) | 2033075 to 8975589 million USA dollars (441%) | 121 |
| 1990-2017 | International migrants | 152.5 to 257.7 million (169%) | 122 |
| 1992-2018 | Total ship number | 28666 to 98755 ships (344%) | 123; Jean Tournadre (in litt., 2020) |
| 1997-2017 | Global merchant fleet tonnage | ~ 723 to ~ 1765 million DWT (244%) | 124 |

<sup>\$</sup>Calculated as percentage increase by dividing the later estimate by the earlier estimate and multiplying by 100. The numbers and percentages given in this Table are approximations whenever numbers are given with a ~, < or > sign because these numbers were taken from graphs in Steffen et al.<sup>2</sup> and ISL<sup>124</sup>.
Abbreviations: DWT = deadweight tonnage is a measure of how much weight a ship can carry; km = kilometre; MT = metric ton; TEU = twenty-foot equivalent unit, a standard size for 6.1 m long containers.

### Increase in the number of outbreaks

From the 1940s to around 2010, the annual total outbreak number (Fig. 1D), the annual total disease number (Fig. 1E), and the annual total country number (Fig. 1F) increased exponentially (Supplementary Data S1). For the annual total country number, this trend continued unabated up to 2020. However, for the annual total disease number and the annual total disease number, the trend reversed since about 2015. While there is some annual up-and-down variation, the overall trends are well demonstrated by the smooth line regressions.

When we control for the number of the global human population, the growing trend for the annual total outbreak number (Fig. 2D) reverses downward around 2010, and the growing trend for annual total disease number (Fig. 2E) actually decreases shortly after 2000 and dramatically in 2020. However, the growing trend for the annual total country number (Fig. 2F) remains the same, which means it continues to increase up to 2020.

A global map of the total outbreak numbers summarized from 1940 to 2020 reveals a highly uneven, positively skewed distribution of outbreaks among countries (Fig. 3) (Supplementary Data S2). The countries with > 250 outbreaks are the USA (1930), the United Kingdom (UK) (810), India (670), Canada (577), Japan (550), Australia (478), China (443), Spain (426), France (406), Italy (381), Germany (377), Brazil (381), and Russia (324).

**Fig. 3.**
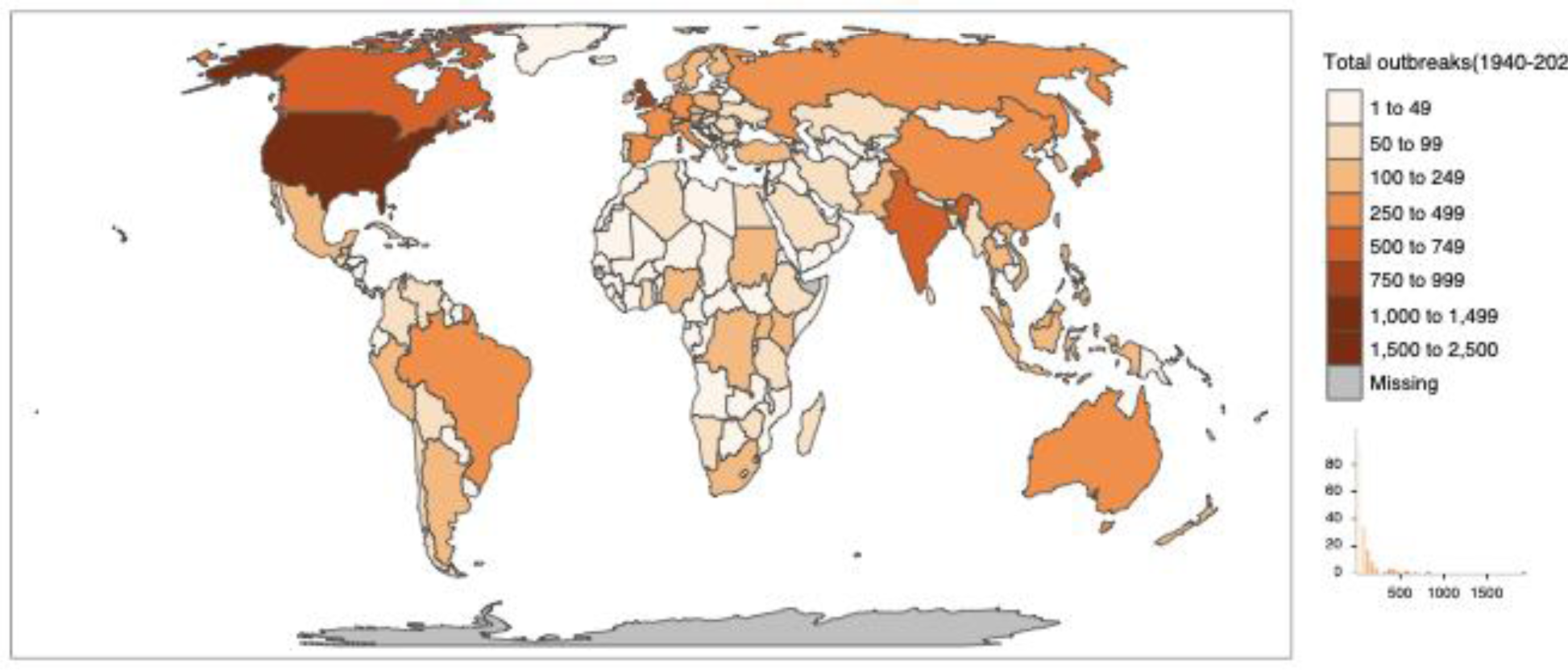
World map of total disease outbreaks from 1940 to 2020. The frequency histogram is also included.

### Decrease in the modularity of networks of countries versus outbreaks

Since the 1940s, the annual modularity measure (which is a measure of the connectivity of the outbreak networks during each year) has overall decreased (Fig. 4). However, it slightly decreased until 1965; after 1965, there is a stronger downward and almost linear trend (Spearman rank test between predicted values by segmented and observed values, rho^2^ = 0.79, F1,77 =297.09, p < 0.00000001). However, in 2020, the value of modularity increased dramatically to a value last seen in the 1970s. Our discontinuity analysis of the trend split the trend from 1940 to 2019 into two parts around an estimated break-point in 1965 (with a standard error of 4.7 years) with (1) a slightly negative linear trend from 1940 to 1965 and (2) a significantly negative linear trend from 1965 to 2019. The global increase in the connectivity of disease outbreaks is also demonstrated by the increase in the number of connections between countries (bipartite graphs in Fig. S1A), the changing modularity among countries (bipartite matrices in Fig. S1B), and the increasing number of clusters and the number of nodes within each cluster (unipartite graphs in Fig. S1C).

**Fig. 4.**
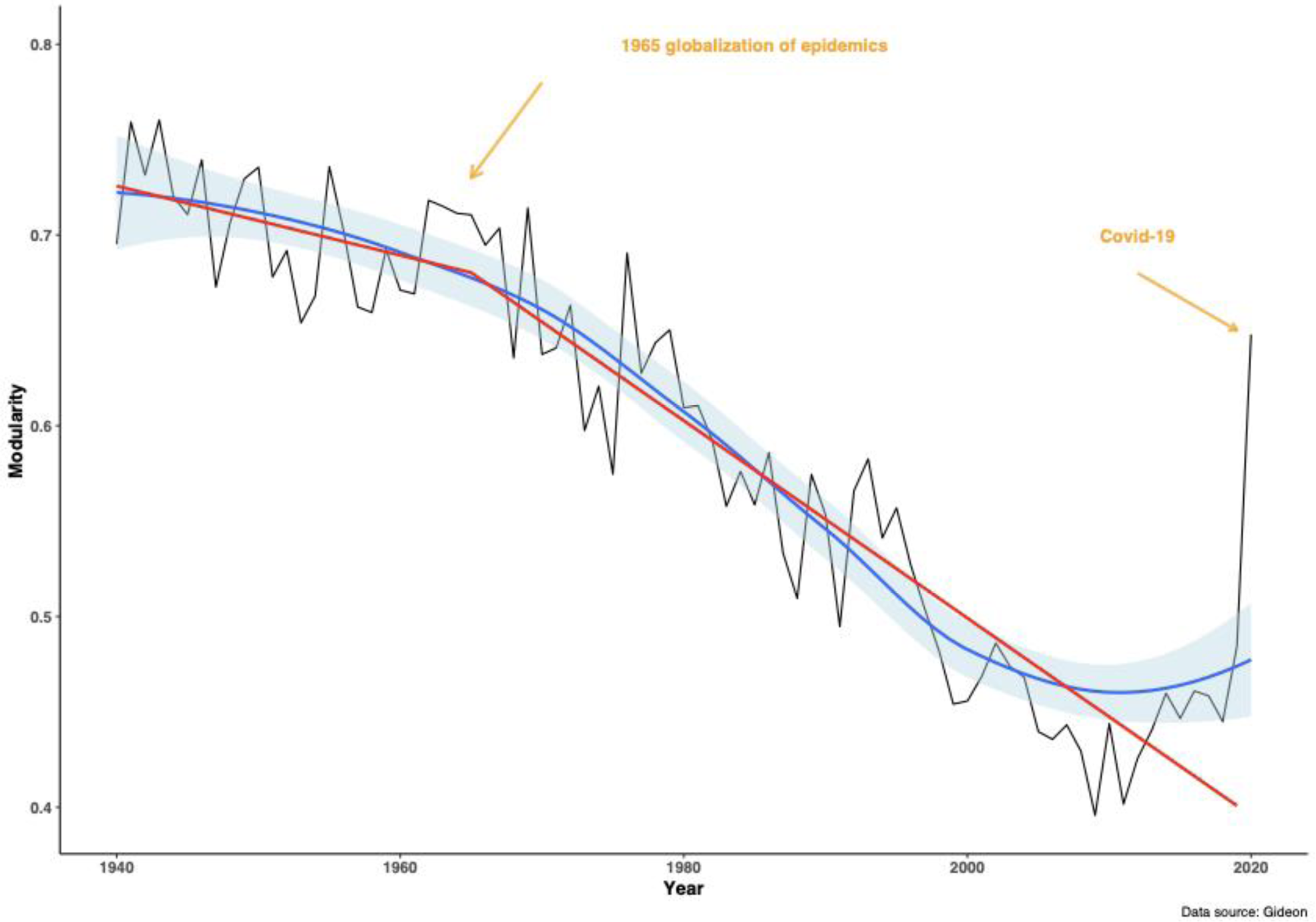
The decrease in the modularity of outbreaks since the 1940s. The graph from plotting the annual modularity measure from 1940 to 2020. We also fitted a smooth regression (in blue) with confidence intervals (in light blue). The two red lines display the results of the discontinuity analysis which split the trend into two parts with an estimated breakpoint in 1965 (see details in Results). The first year of the Covid-19 pandemic resulted in an increase of the modularity measure for 2020 to a level similar to the ones observed in the 1970s.

### Relationship between air travel and modularity

Since the 1970s to 2019, the annual modularity measure was negatively correlated with the annual total number of passengers (Fig. 5A) (Spearman rank test, rho^2^ = 0.80, F1,47 = 190.7, P < 0.00001) and the annual total amount of air freight (Fig. 5B) (Spearman rank test, rho^2^ = 0.78, F1,45 = 164.2, P < 0.00001). However, the addition of the year 2020 to these two graphs demonstrated its outlier position due to the increased value of modularity and the decreased numbers of air passengers and freight in 2020 (Fig. 5A and 5B).

**Fig. 5.**
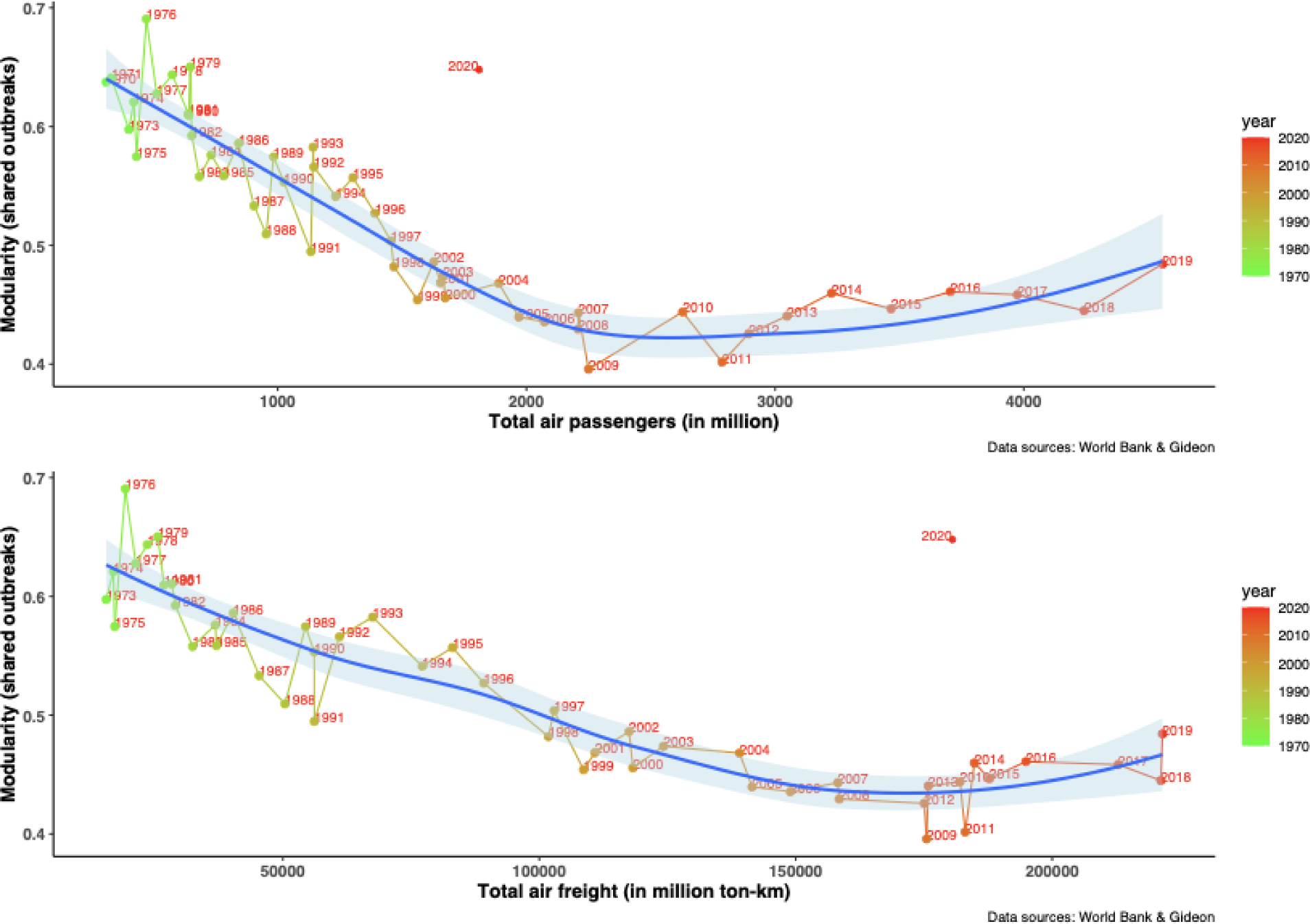
Correlation between annual modularity and in air travel and transport since the 1970s. Resulting graphs from plotting the (A) annual global number of air passengers and (B) annual global total of air freight (cf. Fig. 1) against the annual modularity measure (cf. Fig. 4) from 1970 to 2020. Fitted smooth regressions (in blue) with confidence intervals (in light blue) are shown. The Covid-19 pandemic in 2020 is an outlier in each graph because modularity increased dramatically in 2020 while both the global number of air passengers (A) and annual global total of air freight (B) decreased dramatically.

### Geography of the centrality of outbreaks

We calculated the centrality values for each nation and for each year, and then calculated the mean and standard deviation using all the annual values for each country (Fig. 6) (Supplementary Data S2). The distribution of mean values is positively skewed, with a few nations exhibiting very high centrality values (Fig. 6A). The countries with the highest centrality values of > 0.20 are the USA (0.92), UK (0.60), Japan (0.40), Canada (0.40), Germany (0.32), Australia (0.32), Spain (0.31), India (0.29), Italy (0.29), China (0.28), and France (0.28). The average centrality measure correlated positively with the GDP per capita of the year 2018 with individual countries as data points (Spearman rank test, n = 138 countries, rho^2^ = 0.25, F1,136 = 45,20, P < 0.0000001). The distribution of standard deviation values of centrality is approximately normally distributed, but with countries with high standard deviations concentrated in Asia, Europe, and the Americas (Fig. 6B).

**Fig. 6.**
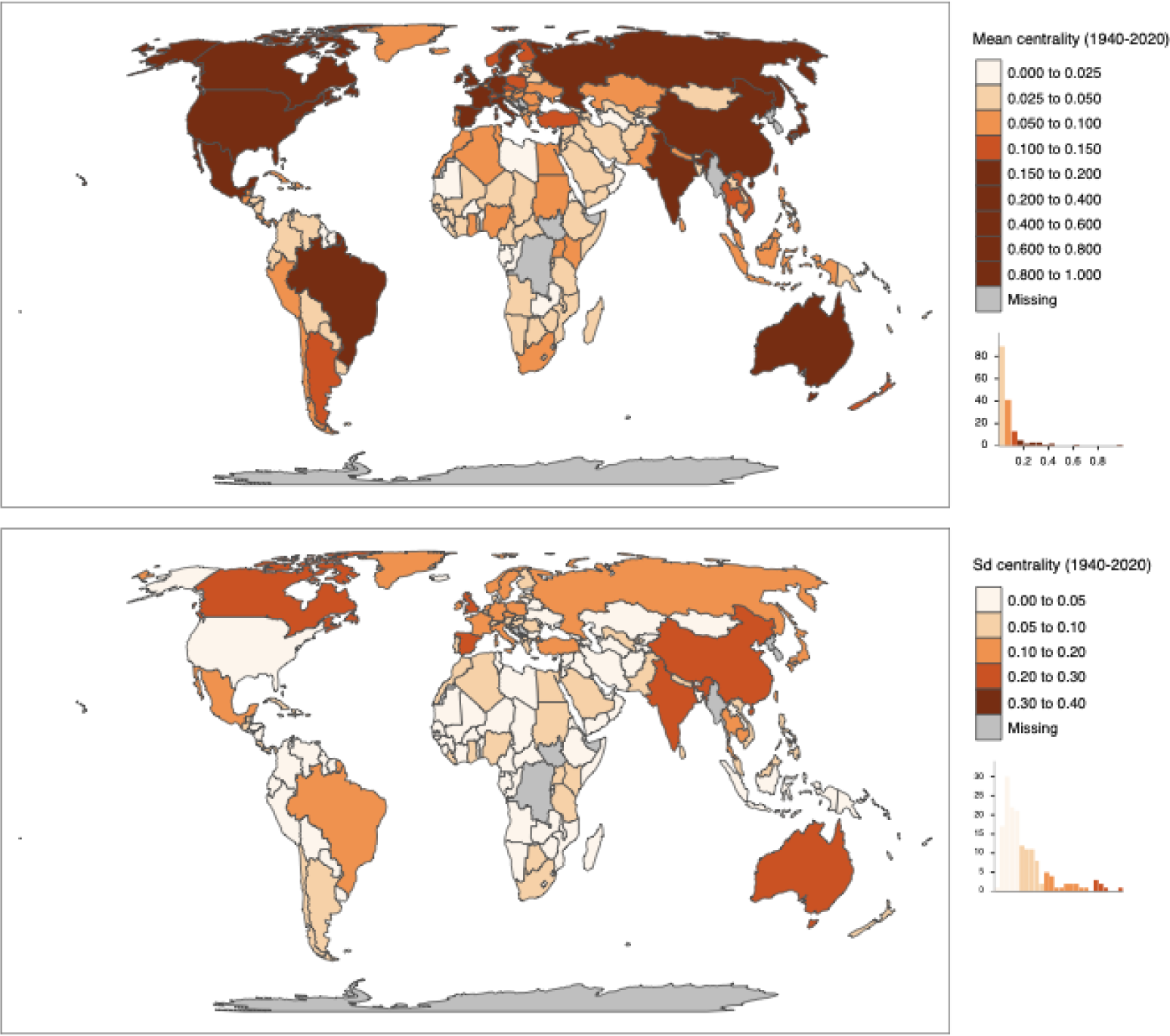
World map of centrality measures from 1940 to 2020. (A) Mean centrality values of shared disease outbreaks and (B) their variability (using the standard deviation). The frequency histograms are also included.

We also plotted all the annual centrality values for each nation within six regions. Despite a large amount of variation over the investigated time period, the three North American countries had the highest mean of centrality values, followed by the three Pacific countries and then the European, South American, Asian, and African countries (Fig. 7). These differences in centrality between the six regions are statistically significant (Kruskal-Wallis χ2 = 869.21, df = 5, P < 2.2e-16).

**Fig. 7.**
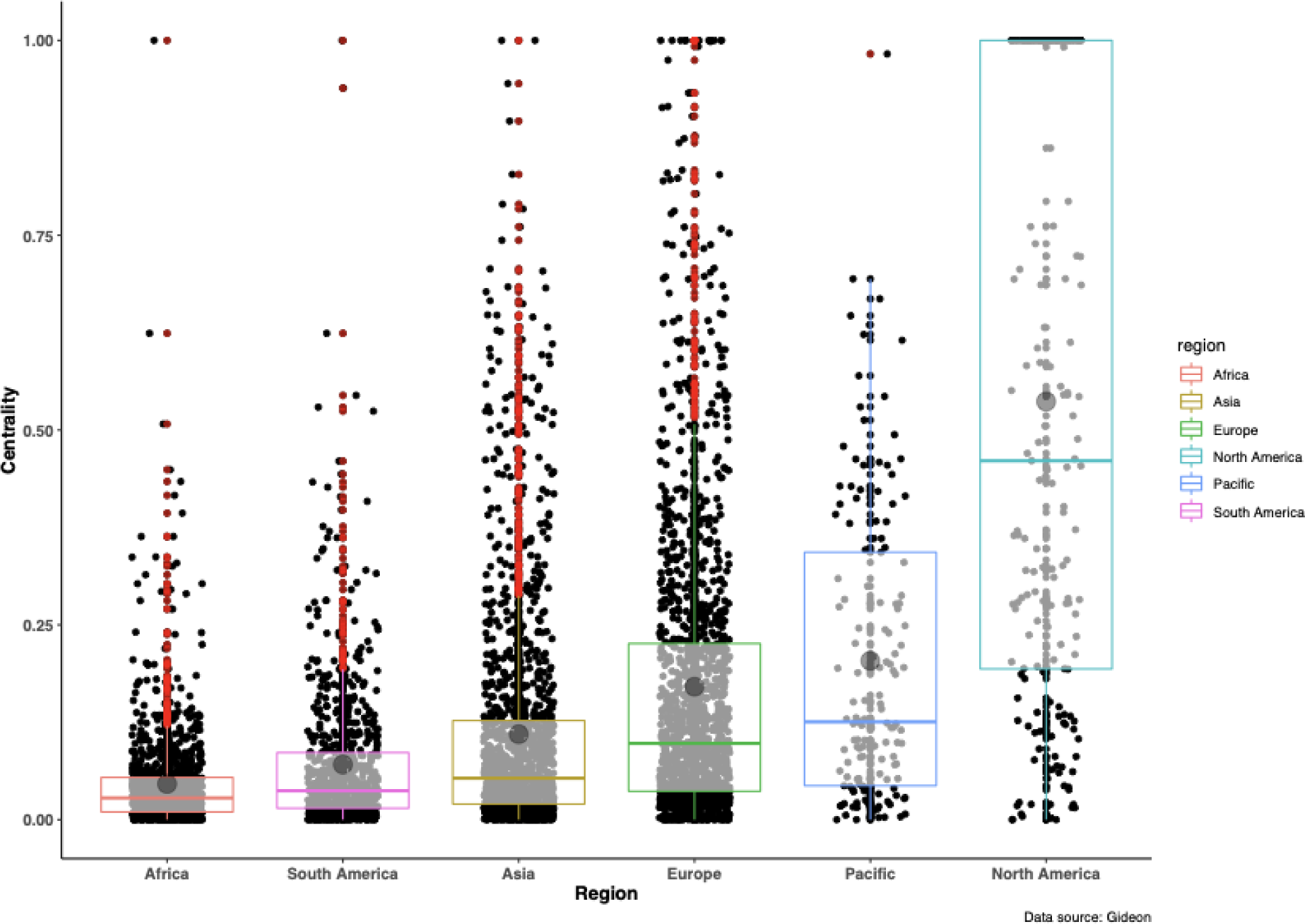
Box and whiskers plots of centrality measures for six regions. Notched boxes of all annual centrality values (1940-2020) for countries in Africa (n = 45 countries), Asia (n = 48 countries), South America (n = 25 countries), Europe (n = 38 countries), Pacific (n = 3 countries), and North America (n = 3 countries). Notched boxes correspond to 95% confidence interval of the interquartile distance between the first and third quartiles (allowing for the comparison of medians), whereby horizontal lines inside the boxes are the medians of the centrality values, grey dots are means of centrality values (with standard deviation represented by vertical lines), and red points are data outliers.

Furthermore, despite the small sample size, the average centrality measure correlated positively with the GDP per capita of the year 2018 with individual regions as data points (Spearman rank test, n = 6 regions, rho^2^ = 0.89, F1,4 = 32.02, P = 0.0048). North American, Pacific, and European countries were more central in our unipartite disease outbreak networks. The most central positions (i.e. the highest centrality values) in the network of all outbreaks were occupied by: the USA for North America; the UK, Germany, and Russia for Europe; Brazil, Argentina, and Cuba for South America; India, Japan, and China for Asia; the Democratic Republic of the Congo, Egypt and Kenya for Africa; and Australia for the Pacific region (Supplementary Data S2).

### Relationship between air connection and centrality

We used the data on flight connections from the OpenFlights85 database to build a network of flight connections among countries. We then calculated the centrality values for each country in this global network of flight connections. The centrality values of 225 countries calculated for the flight connection network correlated with the centrality values of the same countries calculated for the disease outbreak network (Fig. 8A; Spearman rank test, n = 183 countries, rho^2^ = 0.35, F1,181 = 97.06, P < 0.0000001).

**Fig. 8.**
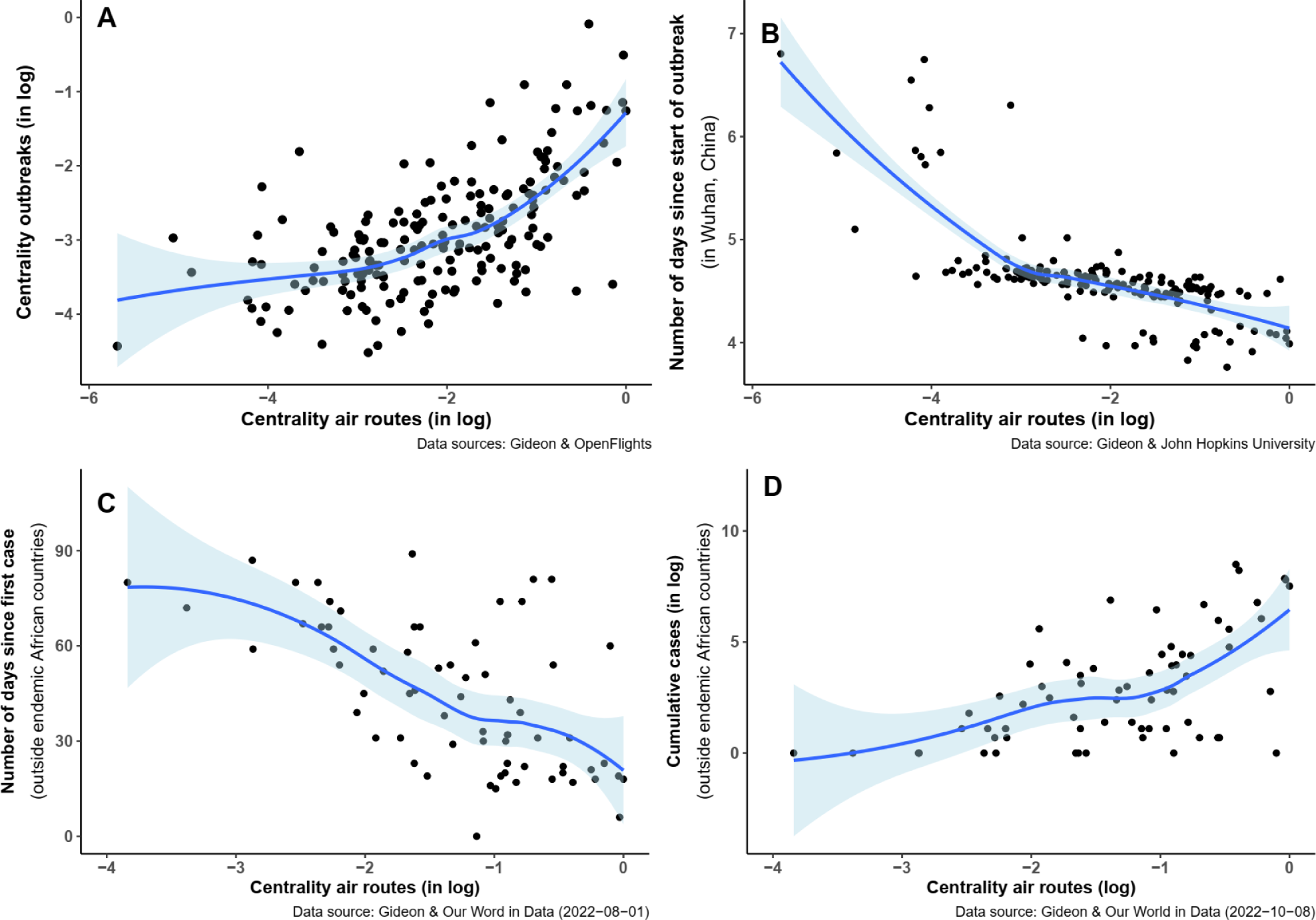
Correlations between centrality values and time of outbreak emergence. Resulting graphs from plotting the centrality values calculated for the flight connection network (A) against the centrality values calculated for the COVID-19 outbreak, (B) the number of days since the start of the COVID-19 outbreak in Wuhan, China, (C) the number of days since the start of the monkeypox outbreak outside of endemic African countries, and (D) the cumulative cases of monkeypox in each country. Fitted smooth regressions (in blue) with confidence intervals (in light blue) are shown.

### Covid-19 and monkeypox outbreaks as test cases

We finally tested our general results with two specific test cases: the outbreak of Covid-19 in 2019 and the outbreak of monkeypox in 2022. We hypothesized that the date of first occurrence of Covid-19 in each country after its emergence in Wuhan (China) at the end of November 2019 should be negatively correlated with the centrality value of each country in the disease outbreak network. The time (in days) between the first occurrence in each country and the start of the outbreak in China correlated negatively with each country’s centrality value (Fig. 8B; Spearman rank test, n = 181 countries, rho^2^ = 0.63, F1,179 = 309.55, P < 0.0000001).

Likewise, we hypothesized that the date of first occurrence of monkeypox in each country after its emergence in Africa in 2022 should be negatively correlated with the centrality value of each country in the disease outbreak network. In this case, we excluded all African countries where monkeypox is endemic, and included all remaining countries. Again, the time (in days) between the first occurrence correlated negatively with each country’s centrality value (Fig. 8C; Spearman rank test, n = 69 countries, rho^2^ = 0.30, F1,67 = 27.45, P < 0.0000001). Moreover, the cumulative cases of monkeypox for each country on 29 July 2022 correlated positively with each country’s centrality value (Fig. 8D; Spearman rank test, n = 69 countries, rho^2^ = 0.34, F1,67 = 34.04, P < 0.0000001).

## Discussion

Our results support the assertion that the Anthropocene is associated with to a great acceleration of infectious disease risks. First, we showed that the number of disease outbreaks, the number of diseases involved in these outbreaks, and the number of countries affected have increased during the entire Anthropocene (Fig. 1D-F). Our results thus expand on the results of previous studies which were more limited in time or space (e.g., references 38, 46, 65-67). Furthermore, these increases have mostly been exponential, although with some recent slowdowns (see Discussion below).

Overall, these trends are even evident when we look at them per capita (Fig. 2D-F), meaning that the global increases (Fig. 1D-F) are not only due to human population growth. Therefore, other factors are likely responsible, such as the various factors mentioned in the Introduction. We note, however, that the per capita trend for the annual total outbreak number slowed down (Fig. 1D) and the per capita trend for the annual total disease number even reversed (Fig. 1E) after 2000. These results mirror results by Smith et al.^38^ who found that per capita cases decreased significantly over time. They suggested that global improvements in prevention, early detection, control and treatment may be responsible.

While the above results only expanded on previously published results, our novel analyses concern the network properties of these outbreaks. Most importantly, we demonstrated that the spatial distribution of these outbreaks has become more globalized in the sense that the overall modularity of the disease networks across the globe has decreased since around 1965 (Figures 4 and S1). In other words, clusters of disease outbreaks began to increasingly become connected with other clusters so that the fragmented nature of outbreak clusters diminished over time. Before 1965, a disease outbreak usually remained confined to one or a few closely connected countries; thereafter, disease outbreaks have become increasingly pandemic in their nature. We thus revealed a long-term, worldwide change in the biogeographic structure of human infectious diseases associated with outbreaks.

We further found that this decrease in modularity is correlated with the increase in air traffic. The increase in global mobility and especially in air traffic (Table 1) allows an outbreak to rapidly spread across several national and continental borders within a short period of time (see also results from modelling and real-world data below).

Third, we demonstrated which countries and regions are most central within these disease networks. Countries which are more centrally located within these disease networks tend to be also the more developed and emerging countries with significantly higher GDPs. Therefore, one cost of increased global mobility is the increased risk of disease outbreaks and their wider spread (although we note that the per capita risk may be slowing down). In further support of these findings, the outbreak of Covid-19 in 2019 and the outbreak of monkeypox in 2022 showed that these outbreaks reached those countries earlier which had a more central location within the global networks. In the case of monkeypox, an earlier arrival also meant that more cases accumulated in the more centrally located countries until the cut-off date of July 2022.

Before we discuss the implications of our results, we address possible limitations and biases. While GIDEON is generally acknowledged to be the most complete, reliable, and up-to-date global dataset on infectious disease outbreaks, we nevertheless should consider that (1) there may have been some underreporting in the early part of the Anthropocene, and (2) recent outbreaks may not have been entered into GIDEON yet.

(1) There may have been some underreporting of infectious disease outbreaks during the early parts of the Anthropocene in developing countries. However, the imposition in the 1980s of the so-called Washington Consensus imposed by the International Monetary Fund, World Bank, and United States Department of the Treasury and the resulting the structural adjustment programs dramatically decreased the public health and education capabilities of many developing countries (e.g., references 68, 69) which should also have affected reporting rates. However, we see no evidence of a slowdown of our trends (Figures 1D-F) in the 1980s and 1990s.

While we cannot exclude the possibility of some underreporting of disease outbreaks during the early part of the Anthropocene, it is rather unlikely that the large increases of several hundreds of percent which we documented in Figures 1D-F are entirely due to underreporting. Since the overall trends are so consistent and so large over a relatively long period of time, we argue that these trends are real even if the actual numbers may be off by a few percentage points.

(2) The recent slowdowns shown in Figures 1D-F could be real, or they could be due to the most recent outbreaks not having been entered into GIDEON yet. GIDEON is updated weekly (see Methods); therefore, the only likely reason for recent underreporting may be that some data might work its way through the system more slowly, e.g., the data reported in national health ministry publications. We note that if the recent decreasing trends in Fig. 1D-F are actually due to underreporting, then this would only make our documented trends even stronger. If they are real, then they are not really influencing the overall decade-long trends very much, although they may be indicative of some recent successes in prevention, control and treatment, as suggested by Smith et al.^38^.

Although it is generally acknowledged that correlation does not prove causation, the negative correlation between air travel and modularity specifically (Fig. 5), and the relationship between increased mobility and the wider spread of disease outbreaks (Figures 1, and 4) in general make sense because they are backed up by numerous theoretical and practical studies (see Text S2). Therefore, it is probable that the staggering global increase in mobility during the Anthropocene (Table 1) is mainly responsible for the increase in (1) the number of disease outbreaks, (2) the number of diseases involved in these outbreaks, (3) the number of countries affected, and (4) the decrease of modularity of these disease outbreaks. However, we acknowledge that other factors may be responsible, especially variables which may covary with mobility measures. Further causal analyses are therefore required, but these are beyond the scope of this study.

Given the lack of any antiviral and, until recently, vaccine treatment, the Covid-19 pandemic forced governments to drastically curtail people’s mobility and introduce continent-wide social distancing and lockdowns in order to at least slow its spread by lowering transmission rates. Thus, the governments’ responses to this acute global health emergency actually mirror many of the recommendations which were given by the various studies about the effects of mobility on the spread of outbreaks (Text S2). It should also be noted that social distancing (called avoidance behaviour in animals) and the restriction of people’s mobility and its most extreme form, namely quarantine, were already used long before the advent of the germ theory of disease, and some of these strategies are even used by other species^70-74^.

Given this link between mobility and disease outbreaks, the key question which decision-makers and society at large should ask is how mobility of humans, animals, and goods (Table 1) should develop in the future, and especially in the case of another life-threatening pandemic. In Text S3, we discuss three possibly scenarios, which we briefly summarize below. While there are of course other scenarios, we hope that our tentative arguments will stimulate further discussion.

The first scenario would continue the current ‘business-as-usual’ trend of ever-increasing mobility without consideration of the costs. These costs include not only the accelerated infectious disease risk, but many other detrimental externalities, such as increased animal and plant disease outbreaks, greenhouse gas emissions, other chemical pollution, noise pollution, land-use changes due to energy production and transportation networks, exotic species introductions, and so on. Indeed, the monkeypox outbreak of 2022 happened after the Covid-19 pandemic was over and the subsequent restart of global travelling, and it clearly showed that the rapid spread of newly emerging diseases remains very high.

‘Business-as-usual’ scenarios foresee tremendous increases in mobility: e.g., a 128% increase for international tourist arrivals until 2030, a 240-1209% increase in global maritime traffic until 2050, and a 213% increase in passenger-km until 2050 (further examples in Text S3). While outbreaks of animal and plant diseases may be amenable to a cost-benefit analysis^75^, the Covid-19 pandemic has clearly shown that simple cost-benefit analyses cannot be applied when the lives of millions of people are at stake. Given that another pandemic becomes more likely with increasing rates of emergence and increasing global mobility, a ‘business-as-usual’ scenario is automatically associated with further epidemics and pandemics, possibly killing millions of humans and devastating local and regional economies or even the global economy. Given all the additional detrimental externalities of ever-increasing mobility, such as scenario should be considered as extremely unsustainable unless the various externalities can be redressed.

The second scenario would attempt to slow down or even reverse mobility rates of infected hosts and vectors, but otherwise attempt to maintain the ‘business-as-usual’ path. In the short term at least, this may be the most realistic and agreeable scenario. The Covid-19 pandemic has demonstrated that identifying infected hosts and reducing secondary infection rates caused by these infected hosts appear to be the most successful strategies to achieve elimination of the outbreak (see Text S3). The required measures, such as mass home quarantine, restrictions on travel, expanded testing and contact tracing, and additional surveillance measures, are thus mostly focused on (1) identifying and isolating infected hosts (which means to drastically restrict their mobility) and (2) drastic restrictions of mobility for uninfected hosts.

However, these restrictions to mobility are hampering economic activity and restrain individual freedoms, so they cannot be a long-term solution. One possible solution is much improved identification and isolation of infected hosts. Therefore, efficient and reliable health checks which can identify various diseases and which can be administered relatively time- and cost-efficiently to large numbers of travellers could be implemented, we may be able to significantly restrict the mobility of infected hosts. Examples of possible measures to be taken include rapid diagnostic techniques, thorough hygienic measures, and much better vector control (examples in Text S3). Such measures could help to decrease the mobility of infected hosts and transmission from them and thus the transmission and global spread of diseases.

The third scenario would intentionally slow down or even reverse mobility rates of humans and other carriers and vectors as part of a much more fundamental reorientation of the global economy along the path of economic degrowth. Thus, decreasing mobility would be part and parcel of a much larger movement towards global sustainability. Declines in mobility would not only decrease the likelihood of epidemic outbreaks, but also decrease the other detrimental externalities and thus reduce humanity’s environmental footprint.

This scenario would likely be opposed on the grounds that, so far, continuous economic growth has almost always been positively linked with increased mobility (see Text S3). However, the many environmental benefits of economic degrowth and deglobalization would be augmented by a global health benefit, the almost certain decrease of infectious disease outbreaks. Since economic degrowth and deglobalization have been advocated by many sustainability experts to deal with the converging environmental crises^19,76-79^, our results further strengthen the argument for such a ‘not-business-as-usual’ scenario. Moreover, the economic degrowth scenario would also ameliorate many of the local conditions or factors associated with the emergence of outbreaks, thus further decreasing the likelihood of disease outbreaks. Finally, the presence of abundant biodiversity and healthy ecosystems has an overall positive effect on human well-being and health^41,49,80-82^ which should count as an additional health benefit of the economic degrowth scenario.

Naturally, decreasing mobility is a moral and political choice which can be informed by science, but not answered by science. However, given all the negative impacts of high mobility, maybe it is time to ask whether it is morally justified, for example, to move the equivalent of all the inhabitants of a small town across a continent so that a football team can be supported by its fans during an away game? Is it necessary to fly halfway around the world for a weekend shopping trip? Is it really most cost-efficient for supply chains to cover the entire globe considering all the resulting externalities? Is long-term sustainability achievable with ever higher rates of mobility?

The demand for ever-increasing mobility is putting many tremendous and increasingly unsustainable stresses on the Earth system and therefore also on many aspects of human well-being and health. In this study, we documented another one: The public health risks of an increasing number of disease outbreaks and their increasingly global spread. Even without the devastating impacts of the Covid-19 pandemic, the additional disease outbreak burden associated with our highly mobile and connected human societies is a definite cost which must be considered in its moral implications as we consider the future trajectory of the Anthropocene^3,10,20^.

## Methods

### Data on global mobility (trade, traffic, transport, travel, and tourism) and GDP

As measures of mobility, we searched for various numerical examples of the increase in global trade, traffic, transport, travel, and tourism during the Anthropocene. The data on the air freight and air passengers were taken from the World Bank database^83,84^. The database yielded the number of air passengers and the air freight in million metric tons (MT) per km from 1970 to 2017 separately for each country. Data on flight connections was downloaded from the OpenFlights^85^ data page. The data on container movements were provided by Container Trades Statistics and the Institute of Shipping Economics and Logistics (in litt., 2020), and the data on total ship numbers by J. Tournadre (in litt., 2020). Other examples were found through a literature search using the appropriate keywords (global migration, mobility, tourism, trade, transport, travel); however, the illustrative examples resulting from this literature search (presented in Table 1) were not intended to be a comprehensive literature review on this topic.

The data on gross domestic product (GDP) per capita was also taken from the World Bank database (https://www.worldbank.org/). GDP per capita was calculated by dividing a country’s GDP by its midyear population; the numbers are in current United States of America (USA) dollars from 1961 to 2018.

To determine trends controlled for global human population (Fig. 2), numbers shown in Fig. 1 were simply divided by the global human population for each respective year.

### Data on infectious disease outbreaks

To collate the total number of infectious disease outbreaks over the years 1940-2018, we extracted the relevant data from the subscription- and web-based global infectious diseases database called Global Infectious Diseases and Epidemiology Network (GIDEON) database^51^ which provides extensive geographical and epidemiological information including outbreaks for 252 recognized infectious diseases in 227 countries and regions. Developed over the past 30 years^86^, GIDEON contains detailed information on the occurrence of epidemic outbreaks of human infectious diseases listed by country and year, as well as the number of surveys conducted in each country.

The GIDEON data are updated weekly with records of confirmed outbreaks. To be as complete as possible, various sources are continually checked for possible inclusion. These sources are all peer-reviewed and backed by scientific evidence^87^ and include the following: (1) ProMED (https://promedmail.org/), the largest publicly-available surveillance system conducting global reporting of infectious diseases outbreaks which provides up-to-date information on outbreaks of diseases which affect humans, animals, and crops grown for food whereby all information is verified by expert teams; (2) the standard publications of the US Centers for Disease Control and Prevention (www.cdc.gov) (3) the Disease Outbreak News of the World Health Organization (https://www.who.int/csr/don/en/); (4) the Outbreaks and Emergencies Bulletin of the World Health Organization (https://www.afro.who.int/health-topics/disease-outbreaks/outbreaks-and-other-emergencies-updates) (5) other dedicated internet lists (e.g., for anthrax and rabies); (6) all available national health ministry publications, both the print and electronic versions; (7) military agencies; (8) standard epidemiological textbooks; (9) data studies presented at major conferences; (10) all relevant peer-reviewed journals which are continually examined for relevant studies, e.g., a monthly search of National Library of Medicine’s Medline (https://www.nlm.nih.gov/medline/) is conducted against a list of all GIDEON key words. Because of its comprehensive and extensive coverage, it has regularly been called “the world’s premier global infectious diseases database”^88^ and “the most comprehensive infectious disease occurrence database currently available at a global scale”^89^.

Therefore, GIDEON is generally considered to be the most complete, reliable, and up-to-date database in the world. To further verify its completeness, Yang et al.^90^ tested GIDEON’s data versus an independently derived dataset for 10 randomly chosen water-associated diseases and found a significant agreement between the two datasets (P < 0001). Furthermore, Talisuna et al.^91^ recently demonstrated the relevance of the GIDEON database for an analysis of the spatial and temporal distribution of infectious disease outbreaks in Africa.

Because of its quality, GIDEON has been used repeatedly in previous macro-scale studies of infectious diseases, epidemics, and pathogen diversity, e.g., for the global distribution of human infectious diseases^88,89,92-94^, of flaviviruses^95^, of leishmaniases^96^, and of water-associated infectious diseases^90^, and for numerous other global analyses (e.g., references 38, 46, 57, 65, 67, 97, 98).

Naturally, the publications and reports included for each country depend on the scientific scrutiny and public health surveillance system of each country and region. Therefore, a cross-country comparison using the number of outbreaks should, for example, control for the number of scientific research reports and investment in the public health system as a proportion of GDP (as in, e.g., Morand and Walther^98^). However, the present study did not compare countries but analyzed the spatial and temporal patterns of outbreaks of infectious diseases on a worldwide basis. We assume that an improved global surveillance system would be better at recording the occurrence of any outbreak. It would also help countries to be better prepared for epidemics. Such improvements of surveillance, preparedness, and prevention should flatten the global trend in epidemics. However, such a global indicator of the quality of the public health surveillance and its response system is still lacking, which may be one reason why there is no slowdown of disease outbreaks (Fig. 1D-F; see also Discussion).

In the context of our study, an infectious disease outbreak is defined as the occurrence of cases of disease in excess of what would normally be expected in a defined community, geographical area or season (more details given in Supplementary Text S1). To be entered into GIDEON, the outbreak has to be defined as such in the source literature. Each row in the GIDEON dataset specifies the disease ‘species’, the year, and the country of the outbreak. The ‘annual total outbreak number’ is simply the annual total number of outbreaks regardless of the disease designation and including all countries. The ‘annual total disease number’ uses the same data for each year as the ‘annual total outbreak number’ but then counts only the different infectious diseases which had at least one outbreak in that respective year. The ‘annual total country number’ uses the same data for each year as the ‘annual total outbreak number’ but then counts only the different countries which had at least one outbreak in that respective year. Our entire 1940-2020 dataset contains 18002 outbreaks of 252 human infectious diseases in 224 nations.

We finally tested our general results with two specific test cases, namely data about the global outbreaks of Covid-19 in 2019 and of monkeypox in 2022. The data for the Covid-19 outbreak were obtained from the COVID-19 Data Repository of the Center for Systems Science and Engineering (CSSE) at Johns Hopkins University (https://github.com/CSSEGISandData/COVID-19, reference 99), and the data for the Mpox (monkeypox) outbreak were obtained from OurWorldInData based on the data produced by WHO (World Health Organization) (https://ourworldindata.org/monkeypox, reference 100). These databases yielded the first occurrence of each disease in each country, which we then correlated with the centrality value of each country in our disease network.

### Statistical analyses

Presence-absence matrices (in our case, with rows for countries and disease occurrences in columns) can be used to calculate and visualize modularity in a number of ways. Here, we used bipartite networks, bipartite modules, and unipartite networks for visualization^60,101^. *Bipartite networks.* We built bipartite networks of presence-absence matrices which link countries (as one set of nodes) with all the recorded epidemic outbreaks (as the other set of nodes). We visualized our bipartite networks as **bipartite graphs** with the package *vegan*, version 2.3-0, implemented in R^102^.

#### Bipartite modules

We also visualized our bipartite networks as **bipartite matrices** (or bipartite modules) with the package *vegan*. Boxes are drawn around each module to identify a group of countries and disease outbreaks that are strongly linked together^60^. We used the function *computeModules* of the package *bipartite* implemented in R to compute modules using the modularity algorithm of Dormann & Strau^103^. The resulting boxed-in modules are detected by this modularity algorithm; in our case, the modules identify those countries which are more connected to each other because they share a common set of infectious disease outbreaks which are, however, less or not shared with other countries.

#### Unipartite networks

We then transformed these bipartite networks where separate nodes from countries were connected with nodes of epidemic outbreaks into unipartite networks using the package *tnet*^104^ implemented in R. We visualized unipartite networks as **unipartite graphs** using the function *cluster_louvain* implemented in the package *igraph*^105,106^. This function is based on a multilevel modularity optimization algorithm^107^.

Within a graph, the same colour refers to a group (i.e. a module) of countries which are more connected to each other because they share a common set of infectious disease outbreaks which are, however, less or not shared with other countries.

#### Modularity measure

Modularity measures computed using bipartite networks or unipartite networks of epidemic outbreaks shared among countries allowed us to identify modules of countries that share common epidemic outbreaks in each respective time period (e.g., a year or a decade) (see references in Introduction and references 101, 107). For the modularity measures obtained for bipartite networks or unipartite networks, we used the modularity measure for the unipartite network for each year calculated with the package *igraph* following the algorithm of Clauset et al.^108^. High modularity calculated in this context means that an epidemic remains relatively constrained within a few countries while low modularity means that an epidemic has spread across relatively more countries.

#### Centrality measure

The calculation of the eigenvector centrality of the unipartite network of each respective year allowed us to determine the number of connections which a country has to other countries in terms of epidemic outbreak sharing. Eigenvector centrality is a measure of the degree of the connectedness of a country and those countries connected to it. High centrality calculated in this context means that a country is connected to many countries which are also well-connected^61,62^.

We used network analyses (i.e., the process of investigating interconnected structures through the use of networks and graph theory) to investigate the sharing of disease outbreaks among countries using the GIDEON database and the flight connections among countries using the OpenFlights^85^ database.

#### Discontinuity time trend

A visual examination of the time trend of the annual modularity measure suggested a discontinuous trend over time. To detect such a discontinuity (or breakpoint) in the trend over time, we used the package *segmented*^109,110^ implemented in R. This package allows the identification of one or more discontinuities using the bootstrap method; in other words, the decomposition of a relationship into one of more piece-wise linear relationships and the identification of breaking point(s). The standard errors of the breakpoint estimates were computed with the procedure of Clegg et al.^111^ which is implemented in the package *segmented*.

#### Smooth regression

We used a smooth regression^112,113^ to visualize the relationships between the independent and the dependent variable. Smoothing is a very powerful regression technique used across all kinds of data analyses. It is designed to detect trends in the presence of noisy data in cases in which the shape of the trend is unknown. In smooth regression, linear regression terms are extended to smooth functions whose exact form is not pre-specified but chosen from a flexible family by the fitting procedures. The underlying assumption behind this technique is that the trend is smooth while the noise is unpredictably wobbly, and no model (or specification of a function) is assumed beforehand (e.g., a linear model). Using only the information contained in the data, this technique can model any functional relationship which is differentiable and smooth. The resulting regression line lets important patterns clearly stand out by eliminating the noise in the data. We used the LOESS (locally weighted smoothing) function implemented in the package ggplot2 implemented in R.

#### Nonparametric statistics

For non-parametric tests, we used the two-sided Kruskal-Wallis test or the Spearman rank test. However, if it was not possible to compute a P-value due to a high number of ties in one variable in consideration, we used the Pearson’s product-moment correlation instead of the Spearman rank test. For these tests, we used the package Hmisc implemented in R.

## Data availability

The GIDEON data is available commercially. We provide four Supplementary Data files (S1 provides data on a yearly basis from 1940 to 2020; S2 provides data on a by-country basis for 203 countries; S3 provides data for the first occurrence of Covid-19 for each country; S4 provides data for centrality values for each country and year; S5 provides data for centrality values and first occurrence of monkeypox; S6 is an R script for producing the figures).

## Code availability

The R scripts for analysing the data and drawing the figures are available from S.M. upon reasonable request.

## Supporting information

Detailed definitions of infectious disease outbreaks

## Acknowledgements

This work was part of theFutureHealthSEA project funded by the French ANR (ANR-17-CE35-0003-01). S.M. is supported by the Thailand International Cooperation Agency (TICA) “Animal Innovative Health”. We sincerely thank Dieter Stockmann from the Institute of Shipping Economics and Logistics (ISL) and Claire Thackeray from Container Trades Statistics (CTS) for providing data about container movements, Jean Tournadre for providing data about ship numbers, and Ting-Wu Chuang for providing references.

## Author contributions

S.M. collated and analysed the data; S.M. and B.A.W devised the study, prepared the figures and tables, and wrote the manuscript.

## Additional information

**Supplementary information** is available for this paper at …

**Competing financial interests:** The authors declare no competing financial interests.

**Correspondence and requests for materials** should be addressed to S.M. and B.A.W.

