## Supplementary material for "The accelerated infectious disease risk in the Anthropocene: more outbreaks and wider global spread": Detailed definitions of infectious disease outbreaks

### The accelerated infectious disease risk in the Anthropocene: more outbreaks and wider global spread

Serge Morand and Bruno A. Walther

**Text S1.** Detailed definitions of infectious disease outbreaks (included in our analyses in the main text), and zoonotic disease outbreaks and emerging infectious disease events (not included in our analyses in the main text).

**Infectious disease outbreaks**

The WHO[1] defined a disease outbreak as follows: "A disease outbreak is the occurrence of cases of disease in excess of what would normally be expected in a defined community, geographical area or season. An outbreak may occur in a restricted geographical area, or may extend over several countries. It may last for a few days or weeks, or for several years."

This definition is also used for entry into the GIDEON database[2], and therefore we follow their definition. Furthermore, GIDEON[3] under the question "How are outbreaks defined in GIDEON?" states that the designation "outbreak" may appear for one of four reasons:

1. An event is specifically [sic] reported as "an outbreak" in source literature.
2. In general, any grouping of cases – including family clusters and epidemics – will be listed as an "outbreak" for purpose of consistency. The term "outbreak" is generic here, and much will depend on the nature of the disease itself as there is no numerical cutoff.
3. Citations of animal disease are denoted as "outbreaks" – even when only one animal is involved – in keeping with OIE [i.e., World Organisation for Animal Health] definitions. Thus, a report of anthrax in a single goat is considered an outbreak in their reporting system.

**Zoonotic disease outbreaks**

The WHO[4] defined a zoonosis as follows: "A zoonosis is any disease or infection that is naturally transmissible from vertebrate animals to humans. Animals thus play an essential role in maintaining zoonotic infections in nature. Zoonoses may be bacterial, viral, or parasitic, or may involve unconventional agents. As well as being a public health problem, many of the major zoonotic diseases prevent the efficient production of food of animal origin and create obstacles to international trade in animal products."

This definition is also used for entry into the GIDEON database[3], and therefore we follow this definition.

**Emerging infectious disease events**

Jones et al.[5] defined emerging infectious disease events as follows: "Here we define the first temporal origination of an EID (that is, the original case or cluster of cases representing an infectious disease emerging in human populations for the first time …) as an EID 'event'."

We are aware that different authors and researchers may use different definitions of infectious diseases and emerging infectious diseases than the ones used in GIDEON[3] and presented above. However, a discussion about the pros and cons of different definitions is beyond the scope of our study.

**Text S2.** A short summary of studies of theoretical models and real-world examples which demonstrated that increased mobility can lead to the faster and wider spread of disease outbreaks.

The emergence of an infectious disease outbreak is due to various local conditions and factors (see Introduction). After emergence, the local, regional, or global spread of a disease is dependent on many other factors of which host mobility is usually one of the most important ones. This is of course especially true for directly transmitted human pathogens (e.g., reference 1 and studies cited below), although mobility of humans as well as vectors is also important for the global spread of vector-borne diseases[2-4].

**Theoretical models** predict that increased mobility leads to a faster and more wide-ranging spread of a disease outbreak, and, vice versa, decreased mobility slows and contains the spread of an outbreak. Modelling the spread of the SARS-CoV-1 pandemic, Hufnagel et al.[5] demonstrated that two control strategies, shutting down airport connections and isolating cities, reduced the spread of the virus. Drastic travel limitations also delayed a pandemic by a few weeks in a model of the global spread of influenza[6]. Similarly, increasing levels of (1) isolation of infectious hosts, household quarantine, and related behavioral changes which reduce transmission rates and (2) air traffic reduction increasingly slowed the global spread of influenza, although the latter control strategy required the almost complete halt of global air traffic in order to have a noticeable effect[7-12]. Epstein et al.[11] also emphasized that a combination of both control strategies would be even more effective, a result mirrored by Mao[13] for a model at the city scale. Crucially, Hollingsworth et al.[10] also showed a significant decrease in the number of countries affected if travel reductions are combined with other control strategies to reduce transmission rates. This result was confirmed by Cooper et al.[7] and Flahault et al.[9] who found that fewer cities (distributed around the world) were affected by a major outbreak if sufficiently early and significant travel and transmission reductions were implemented. In a simulated smallpox attack, even gradual and mild behavioral changes had a dramatic impact in slowing an epidemic[14]. Another model suggested that public health policies that encourage self-quarantine by infected people can lower disease prevalence[15].

**Real-world examples** also demonstrate the link between increased mobility and faster disease outbreaks. Real data on influenza in the USA showed that a reduction in air travel resulted in a delayed and prolonged influenza season[16,17]. Similarly, the presence of airports and railway stations significantly advanced the arrival of influenza during the 2009 pandemic in China[18]. Global connectivity due to air traffic allows an outbreak to rapidly spread across several national and continental borders within a short period of time. For example, the 2014 Ebola virus outbreak spread from Africa to several continents via international air travel[19]. Sevilla[20] reviewed and modelled how air travel can aid the global spread of Ebola, H1N1 influenza, SARS-CoV-1, and pneumonic plague. A systematic review of the effectiveness of travel reductions concluded that internal travel restrictions as well as international border restrictions both delay the spread of influenza epidemics[21].

**Text S3.** A brief discussion of three possible scenarios how global mobility levels could continue or change due to societal decisions.

Given the link between mobility and disease outbreaks documented by our study, the key question which decision-makers and society at large should ask are which of the following three scenarios should we aim for in the coming decades. We only outline them in very broad terms, because their purpose is to stimulate further discussion, not to be a comprehensive review of all possible scenarios and outlooks.

(1) Once the current SARS-CoV-2 pandemic is over, we continue on our 'business-as-usual' path of ever-increasing mobility without regard to the costs in terms of the accelerated infectious disease risk.

(2) We slow down or even reverse mobility rates of infected hosts and vectors, but otherwise attempt to maintain the 'business-as-usual' path.

(3) We intentionally slow down or even reverse mobility rates of humans and other carriers and vectors (in other words, decrease many or all of the mobility measures in Table 1) as part of a much more fundamental reorientation of the global economy along the path of economic degrowth. Thus, decreasing mobility would be part and parcel of a much larger movement towards global sustainability.

We briefly discuss some of the implications of each scenario. However, this discussion is by no means exhaustive, but meant to stimulate further discussion and study which are urgently needed to deal with the accelerated infectious disease risk of the Anthropocene.

(1) Most likely, at least in the short-term, economic and political decision-makers will return to 'business-as-usual' which means increasing mobility rates even more. After all, various projections predict further tremendous increases of mobility within the next few decades. For example, international tourist arrivals worldwide are expected to increase by 3.3% a year between 2010 and 2030 to reach 1.8 billion by 2030[1] from the 1.4 billion recorded in 2018 (Table 1). WATF[2] predicted a 3.7%, 2.3%, and 2.0% annual increase in passenger traffic, air cargo, and aircraft movements, respectively, until 2040. Sardain et al.[3] predicted a 240-1209% increase in global maritime traffic by 2050 while the ITF[4] projected an average 3.4% annual growth in the demand for global freight until 2050. Over the next 20 years, air passenger traffic and the number of flying air freighters are predicted to increase by 100% and 30%, respectively[5]. Purwanto et al.[6] predicted that, from 2005 to 2050, world passenger transport will increase from 34.2 to 73.0 trillion passenger-km, world freight demand from 26.3 to 50.8 billion MT-km, the world fleet of passenger cars from 735 to 1400 million vehicles, the world fleet of road freight vehicles from 107 to 160 million vehicles, the world fleet of rail vehicles from 297000 to 622000 vehicles, the world fleet of aircraft from 13410 to 21335, and the maritime fleet from 901 to 2027 million dwt.

In addition to the tremendous environmental and social costs and risks of this scenario (e.g., increasing land-use change, greenhouse gas emissions, resource use and waste production, etc.), which includes the risk of widespread ecosystem collapse (see Introduction), we can now add the cost of an increasing infectious disease risk. While our study only focused on human infectious diseases, this cost related to increased mobility is also increasing for animal and plant disease outbreaks as well as alien species introductions (e.g., references 3, 7-10). While outbreaks of animal and plant diseases may be amenable to a cost-benefit analysis[11], the current SARS-CoV-2 pandemic has clearly shown that simple cost-benefit analyses cannot be applied when the lives of millions of people are at stake. Given that another pandemic becomes more likely with increasing rates of emergence and increasing global mobility, a ‘business-as-usual’ scenario is automatically associated with further epidemics and pandemics, possibly killing millions of humans and devastating local and regional economies or even the global economy.

If a ‘business-as-usual’ scenario is followed with regards to global mobility, countries and the world community should at least invest in better detection and surveillance methods to catch and contain the next pandemic as early as possible, and in better preparedness of public health facilities in case the next pandemic nevertheless gets out of hand[12-14]. However, this scenario nevertheless will likely be associated with an increased number of epidemic outbreaks (some of which may become devastating pandemics), given the results of our study.

(2) Consequently, the most realistic and agreeable scenario may be to slow down or even reverse the mobility rates of infected hosts and vectors. The current SARS-CoV-2 pandemic has demonstrated that identifying infected hosts and reducing secondary infection rates caused by these infected hosts appear to be the most successful strategies to achieve elimination of the outbreak (e.g., references 15, 16). The required measures, such as mass home quarantine, restrictions on travel, expanded testing and contact tracing, and additional surveillance measures, are thus mostly focused on (1) identifying and isolating infected hosts (which means to drastically restrict their mobility) and (2) drastic restrictions of mobility for uninfected hosts.

While the latter is possible in crisis situations, it cannot be a long-term solution to the quandary of the increased infectious disease risk of the Anthropocene unless we want to decrease our total global mobility (see scenario 3 below). Therefore, much improved identification and isolation of infected hosts may be the way forward.

Already, such measures have been adopted during the current SARS-CoV-2 pandemic, e.g., body temperature checks for every air travel passenger even though they appear to be rather ineffective and unreliable because many hosts are asymptomatic[17]; rather, hygiene measures, mask use, and distancing were more effective[18]. Later on, mass testing was introduced which can be effective at reducing transmissions if done properly and comprehensively[19,20]. For air travel, testing of every passenger using saliva-based, rapid testing or even detection with the help of ‘sniffer dogs’ was suggested as a way forward[18,21].

Therefore, if efficient and reliable health checks which can identify various diseases and which can be administered relatively time- and cost-efficiently to large numbers of travellers could be implemented, we may be able to significantly restrict the mobility of infected hosts. While such a proposal may sound like “pie in the sky” at the moment, rapid advances in diagnostic techniques, such as translational proteomics, may soon allow us to identify infected hosts using simple breathalyzer, saliva, or urine tests[21-24]. Furthermore, hygienic measures, such as the enforcement of handwashing and the wearing of efficient facemasks in public transport hubs, complete and regular disinfection of important traffic hubs and vehicles (including the air and all surfaces), and much better vector control should become mandatory global standards of public health (e.g., references 25, 26, reviewed in Huizer et al.[27]), especially in the most central of traffic hubs, such as the world's most connected airports[18,28,29]. Flight bans can also be effective, but must be managed carefully[30]. Such measures could help to decrease the mobility of infected hosts and thus the transmission and global spread of diseases if sufficient coverage is achieved.

(3) The most sustainable scenario is, however, to slow down or even reverse global mobility rates of humans and other carriers and vectors, especially if it is part and parcel of a much larger movement towards global sustainability by reducing humanity's environmental footprint and replacing unsustainable economic growth with sustainable economic degrowth[31-38]. Such a general, comprehensive and global slowdown of mobility of both uninfected and infected people and vectors would be opposed for many reasons and by many interest groups, mainly based on economic arguments about the need for continuous economic growth which has so far almost always been positively linked with increased mobility (e.g., references 39-42). It is to some extent possible to decouple mobility from economic growth[43,44], but even if such a decoupling was achieved, it would not sufficiently reduce mobility to decrease significantly infectious disease risks. As the modelling results cited above and the experience with the current SARS-CoV-2 pandemic clearly show, only a huge reduction in mobility and contact rates is sufficient to achieve a slowdown or halt of a highly contagious disease outbreak which is already under way.

Yet, a decrease in global levels of mobility should also decrease the overall number of disease outbreaks as well as their global reach, according to our results (which is different to just slowing and containing the spread of a specific outbreak, see Text S2). Therefore, the many environmental benefits of economic degrowth and deglobalization would be augmented by a global health benefit. Since economic degrowth and deglobalization have been advocated by many sustainability experts to deal with the currently converging environmental crises (climate change, ocean acidification, biodiversity, etc., see references in main text), the results of our study further strengthen the argument for such a 'not-business-as-usual' scenario.

Moreover, the economic degrowth scenario would also ameliorate many of the local conditions or factors associated with the emergence of outbreaks (see Introduction), thus further decreasing the likelihood of disease outbreaks. Finally, we have a growing understanding that the presence of abundant biodiversity and healthy ecosystems has an overall positive effect on human well-being and health[45-49] which should count as an additional health benefit of the economic degrowth scenario.

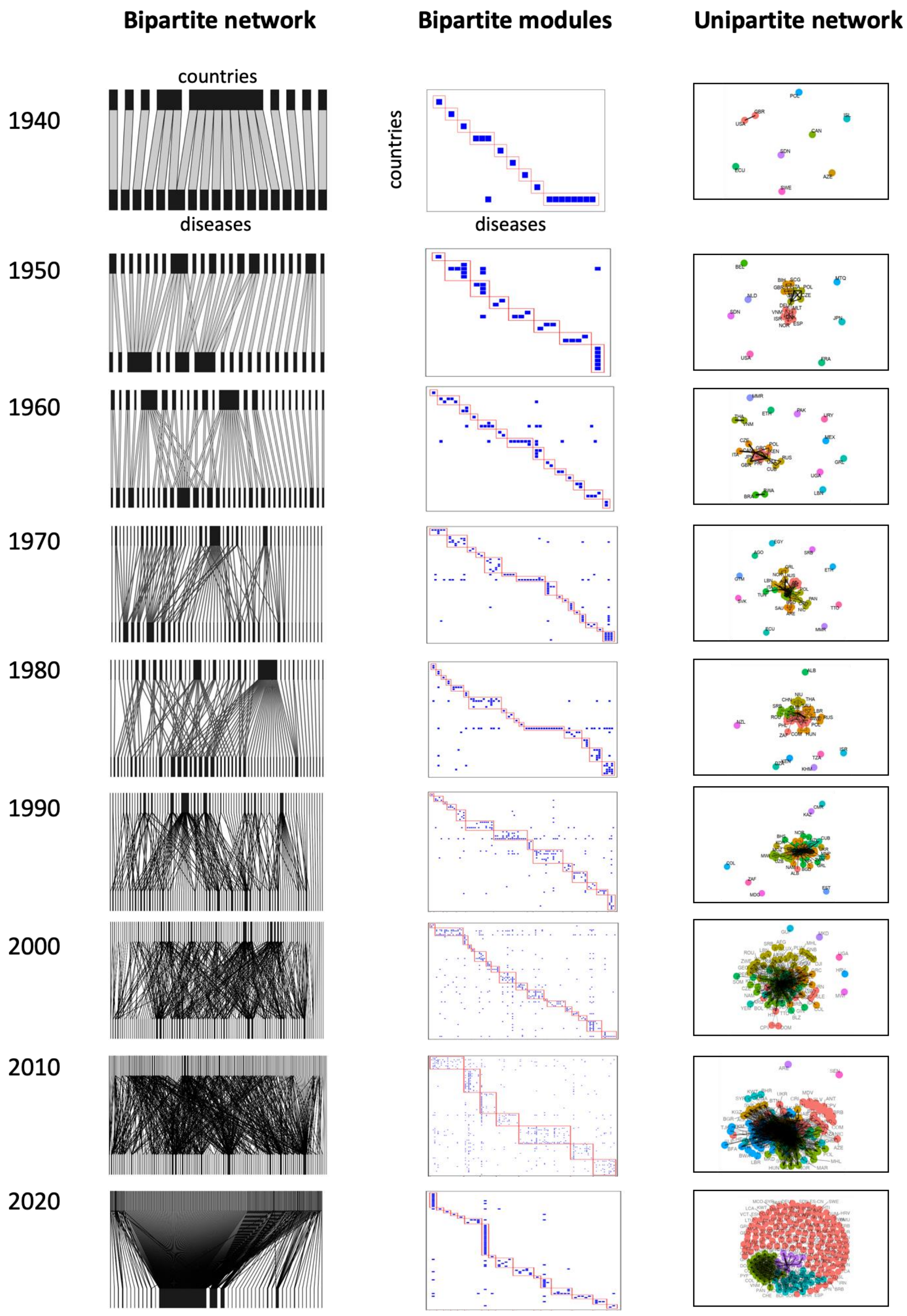


**Fig. S1. | Bipartite graphs, bipartite matrices, and unipartite graphs for the seven decades of the Anthropocene.** Visualizations of the decrease in modularity of disease outbreaks from the 1940s to the 2020s utilizing (A) bipartite graphs, (B) bipartite matrices, and (C) unipartite graphs of disease outbreaks among countries. The plots of the bipartite matrices (B) identify the modules with red boxes. The plots of the unipartite graphs identify all the countries belonging to one module in the same colour, with each module being assigned a different colour.

**Supplementary Data S1. Data on a yearly basis from 1940 to 2020 (data.year.global.xlsx)**

Original data on the first worksheet ‘data.year.global’

Year
Total diseases = total counts of the different infectious diseases which had at least one outbreak per year extracted from GIDEON database
Total outbreaks = total number of outbreaks regardless of the disease designation per year extracted from GIDEON database
Total countries = total number of countries with a least one outbreak in the year from GIDEON database
Word population = total world population per year extracted from World Bank database
Air transport freight (million ton-km) = total world air freight in million metric tons (MT) per km by year extracted from World Bank database
Total passengers = total world number of air passengers per year extracted from World Bank database
GDP (constant 2015 US$) = total world gross domesric product (in contsant 2015 US$) per year extracted from World Bank database
GDP per capita (current US$) = world gross domestic product (GDP) per capita per year extracted from World Bank database
Modularity (outbreaks) = values of the modularity of the unipartite network of shared outbreaks among countries computed for each year from 1940 to 2020

Metadata on the second worksheet

**Supplementary Data S2. Data on a by-country basis for 203 countries (data.country.xlsx)**

Original data on the first worksheet ‘data.country’

Country = country name
Country code = ISO a3 code of the country
GDP per capita (current US$) 2019 = gross domestic product (GDP) per capita for the year 2019 extracted from World Bank database
Total outbreaks = total number of outbreaks regardless of the disease designation from 1940 to 2018 extracted from GIDEON database
Total diseases = total counts of the different infectious diseases which had at least one outbreak from 1940 to 2018 extracted from GIDEON database

Centrality (average) = average values of the eigenvector centrality of the unipartite network of shared outbreaks among countries computed for each year from 1940 to 2018

Standard deviation centrality = standard deviation of the average values of the average centrality

Centrality (airport) = values of the eigenvector centrality of the unipartite network of shared air routes among countries. The values were computed using the data of the year 2014 from OpenFlights

Metadata on the second worksheet

**Supplementary Data S3. Data for the first occurrence of Covid-19 for each country (data.covid.time.xlsx)**

Original data on the first worksheet 'data.covid.time'

Country = name of the country

Date = date of the first occurrence of Covid-19, data from COVID-19 Data Repository of the Center for Systems Science and Engineering 454 (CSSE) at Johns Hopkins University, https://github.com/CSSEGISandData/COVID-19

Metadata on the second worksheet

**Supplementary Data S4. Data for centrality values for each country and year (data.centrality.country.1940.xlsx)**

Original data on the first worksheet 'data.centrality.country.1940'

Country = name of the country

Region = major regions of the world

Year = from 1940 to 2020

Centrality = values of the eigenvector centrality of the unipartite network of shared outbreaks

Metadata on the second worksheet

**Supplementary Data S5. Data for centrality values and first occurrence of monkeypox (data.mpox.country.xlsx)**

Original data on the first worksheet 'data.mpox.country'

Country = country name

Country code = ISO a3 code of the country

Centrality (airport) = values of the eigenvector centrality of the unipartite network of shared air routes among countries. The values were computed using the data of the year 2014 from OpenFlights

Begin monkeypox = date of the first case of monkeypox (Mpox) reported in the country from WHO, compiled in Our Word in Data https://ourworldindata.org/monkeypox

Total number of cases = total number of cases of monkeypox (Mpox) at the date of 2022-08-01 reported in the country from WHO, compiled in Our World in Data https://ourworldindata.org/monkeypox

Metadata on the second worksheet

**Supplementary Data S6. R script for producing the figures (gideonPlot.R)**
